# RNA-modifying enzymes shape virulence and stress response in *Staphylococcus aureus*

**DOI:** 10.64898/2026.11.25.769361

**Authors:** Chloé Teixeira, Florence Couzon, Pauline François, François Vandenesch, Stefano Marzi, Karen Moreau

**Affiliations:** CIRI, Lyon 1 Université, INSERM U1111, CNRS UMR5308, ENS de Lyon, Lyon, France; Centre National de Référence des Staphylocoques, Institut des agents infectieux, Hospices Civils de Lyon, Lyon, France; Université de Strasbourg, CNRS, Architecture et Réactivité de l’ARN, 67000 Strasbourg, France

## Abstract

In addition to transcriptional regulation, RNA modifications can affect bacterial physiology by modulating translation efficiency and protein production. This study investigated the conservation of genes predicted to participate in tRNA and rRNA modification pathways in *Staphylococcus aureus* (*S. aureus*) and screened selected tRNA- and rRNA-modifying enzyme transposon mutants for associated phenotypes. A panel of 52 candidate loci was initially identified in the reference strain USA300_FPR3757, and their functional annotations and predicted RNA targets were subsequently curated using recent species-specific RNA modification maps. Comparative analysis of 301 clinically diverse strains showed that most candidate loci were highly conserved, defining a broadly conserved repertoire of RNA-modification pathways in *S. aureus*. Selected tRNA- and rRNA-modifying enzyme transposon mutants were then examined for alterations in growth, responses to antibiotic and oxidative stress, host-cell interactions, and production of virulence-associated proteins. Most mutants showed no major growth defect in rich medium, whereas several displayed reduced growths in RPMI medium. The transposon insertions had limited effects on antibiotic susceptibility, although mutants carrying insertions in *truA*, *miaA* or *miaB* showed altered survival under antibiotic exposure. The *miaB* transposon mutant also exhibited reduced survival under oxidative stress, while mutants carrying insertions in *rlmL*, *rlmN*, *miaB* or *tadA* displayed reduced host-cell adhesion and/or macrophage uptake. Finally, several mutants showed altered levels of toxins or immune-evasion proteins. Overall, this phenotypic screen identifies associations between RNA-modification pathways and condition-specific traits related to stress adaptation and host interactions in *S. aureus* and provides a basis for future studies.

**IMPORTANCE:** RNA modifications are increasingly recognized as contributors to bacterial adaptation, but their roles in the physiology and pathogenicity of Gram-positive bacteria remain poorly understood. Here, we examined the conservation of predicted tRNA- and rRNA-modifying enzymes across clinically diverse *Staphylococcus aureus* (*S. aureus*) strains and performed a broad phenotypic screen using a collection of transposon mutants. Several mutants displayed condition-specific alterations in antibiotic and oxidative-stress responses, interactions with host cells, or production of virulence-associated proteins. These results identify RNA-modification pathways potentially involved in the adaptation of *S. aureus* to conditions encountered during infection and the fine-tuning of virulence. They also provide a basis for future mechanistic studies to determine how individual RNA modifications contribute to these phenotypes.

## INTRODUCTION

*Staphylococcus aureus* (*S. aureus*) is a Gram-positive bacterium and a major human pathogen; it is among the leading causes of mortality associated with bacterial infections worldwide, ranking second only to *Mycobacterium tuberculosis* according to the latest global estimates (1). *S. aureus* causes infections ranging from superficial skin and soft-tissue infections to severe invasive diseases, including pneumonia, endocarditis, osteomyelitis and bloodstream infections (2). Its pathogenicity relies on a diverse repertoire of virulence factors that promote adhesion to host cells, tissue invasion, immune evasion and adaptation to environmental and host stresses (3–5). The expression of these factors is finely controlled by interconnected regulatory systems, including transcription factors, regulatory RNAs or two-component systems that respond to environmental and host-derived signals (6).

RNA modification represents an additional mechanism capable of modulating gene expression and cellular physiology. RNA modifications are introduced by specific enzymes and occur in all major classes of RNA. By influencing RNA folding, stability, molecular interactions and decoding, these chemical modifications can affect translation efficiency, fidelity and protein production (7). More than 400 distinct RNA modifications are currently listed in the MODOMICS database (8). They range from relatively simple chemical alterations, including base or ribose methylation, isomerization, reduction, thiolation and acetylation, to hypermodified nucleotides generated through multistep biosynthetic pathways. For example, queuosine synthesis and incorporation require the coordinated activity of several proteins (9, 10). Many RNA-modifying enzymes and pathways are conserved among bacteria, although their substrates and physiological functions can vary between species. Most enzymes preferentially modify a particular class of RNA, but some display broader substrate specificity. For example, while TadA is best known for catalyzing the deamination of adenosine at position 34 (A34) to inosine in tRNAᵃʳᵍ(ACG), its ability to edit specific messenger RNAs (mRNAs) has also been documented in several bacterial species (11, 12). In *Escherichia coli* (*E. coli*), RlmN catalyzes the formation of m^2^A37 in several transfer RNAs (tRNAs) and m^2^A2503 in 23S ribosomal RNA (rRNA) (13). However, this dual specificity does not appear to be conserved under the conditions examined in *S. aureus*: m²A2530 was detected in 23S rRNA, whereas m²A37 was not detected in any analyzed tRNA despite the presence of an RlmN orthologue (14). This illustrates that functional annotations derived from model organisms cannot necessarily be transferred directly to *S. aureus* without species-specific experimental validation.

tRNAs contain the greatest diversity and density of characterized RNA modifications in bacteria, followed by rRNAs, whereas modifications in mRNAs and other non-coding RNAs remain less extensively characterized (8). tRNA modifications have different impact on the translation process depending on their positions. Modifications in the anticodon loop commonly affect decoding efficiency, reading-frame maintenance and translational fidelity, whereas modifications within the tRNA body frequently contribute folding, stability and conformational dynamics (15). rRNA modifications are concentrated in functionally important regions of the ribosome, including the peptidyl-transferase center of the 50S subunit and the decoding center with the tRNA and mRNA binding sites of the 30S subunit (16). Through effects on base pairing, stacking, hydration and local conformational dynamics, pseudouridylation, nucleotide reduction and different forms of methylation can influence rRNA structure, ribosome assembly and translation (17–19). Nevertheless, the consequences of an individual modification depend on its location and structural environment and cannot be inferred solely from its chemical class.

The contributions of RNA modifications to bacterial stress adaptation and pathogenicity have received increasing attention over the past decade, although most studies have focused on Gram-negative bacteria and on individual enzymes, modifications or experimental conditions (15, 19). For example, the effects of disrupting several RNA-modification pathways during antibiotic exposure have been investigated in *Vibrio cholerae* (20). Changes in RNA-modification abundance have also been examined under hypoxia in *Mycobacterium bovis* (21) and in response to temperature variation in members of the order Bacillales (22). By comparison, the collective contribution of tRNA- and rRNA-modification pathways to the physiology and host-associated phenotypes of *S. aureus* remains poorly defined.

In this study, we examined the distribution and conservation of candidate RNA-modification loci in clinically diverse *S. aureus* strains and performed a broad phenotypic screen of selected RNA-modifying enzymes transposon mutants. We evaluated growth in rich and nutrient-limited media, responses to antibiotic and oxidative stress, interactions with host cells, and production of virulence-associated proteins. The screen identified locus- and condition-specific associations between candidate RNA-modification pathways and several phenotypes related to stress adaptation or host interactions. Rather than supporting a single global regulatory role of tRNA and rRNA modifications, the results define a set of candidate pathways for subsequent mechanistic analyses aimed at determining how the observed phenotypes result directly from loss of the corresponding RNA modifications.

## RESULTS

### Most candidate RNA-modification loci are highly conserved across *S. aureus* clinical strains

To compile an initial set of candidate RNA-modification genes in *S. aureus,* we collected enzymes previously characterized in other bacterial species (23–26) or listed in the MODOMICS database (8), and identified their putative homologues in USA300_FPR3757 reference genome. This procedure generated an initial panel of 52 candidate loci that was used for the comparative genomic analyses described below. Following completion of these analyses, the functional annotations, nomenclature and predicted RNA targets were revised using recently available *S. aureus*-specific tRNA and rRNA modification maps (14, 26, 27). The resulting curated inventory is presented in Table 1. Candidates added during this subsequent curation were not included in the original 52-locus comparative analysis.

**Table 1:**
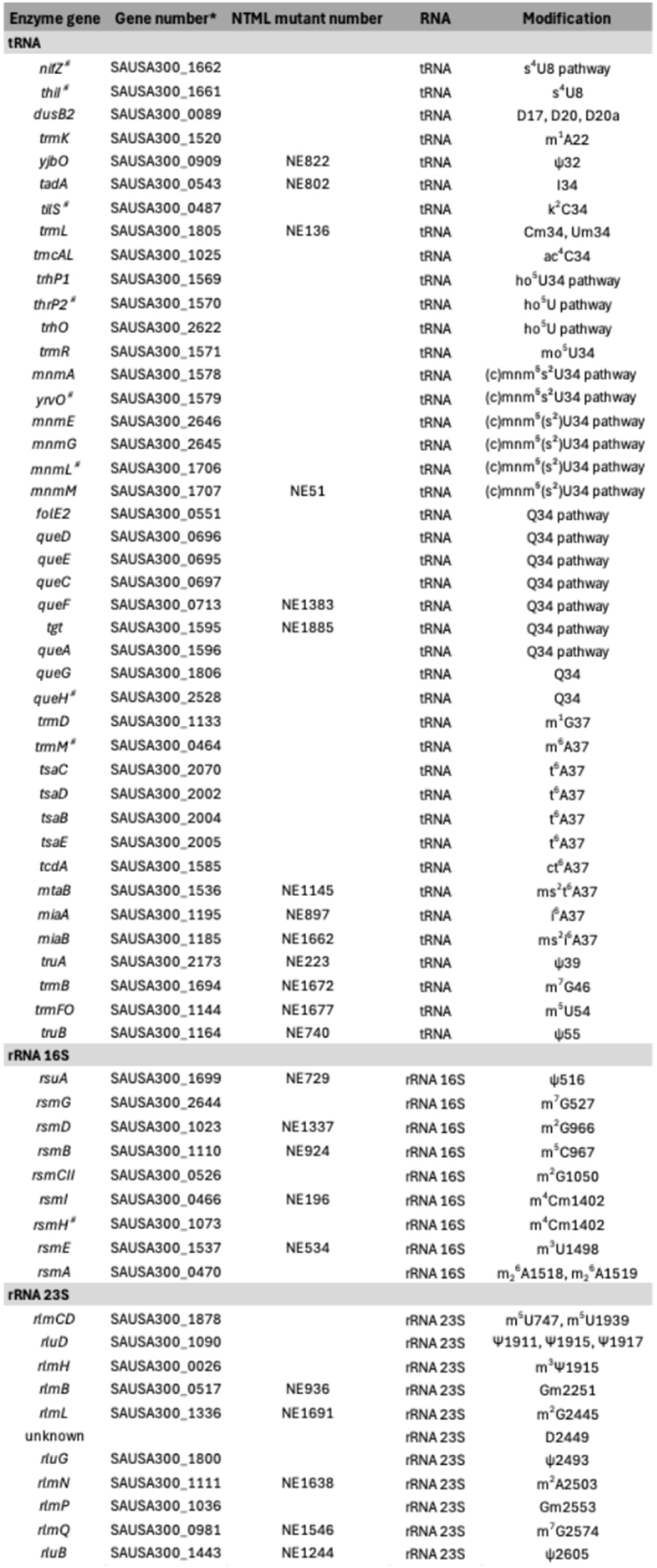
RNA modifying enzymes and associated tRNA- and rRNA-modifications in S. aureus. tRNA modifications enzymes have been defined from Jaramillo-Ponce *etal*. 2026 (14). fRNA modifications enzymea have been defined Banhenea-Cedr0 2023 (27). * fromUSA300_FPR3757 annotation * not included in the initial set of 54 candidate loci

| Enzyme gene | Gene number* | NTML mutant number | RNA | Modification |
|---|---|---|---|---|
| tRNA |  |  |  |  |
| *nifZ*<sup>#</sup> | SAUSA300_1662 |  | tRNA | s<sup>4</sup>U8 pathway |
| *thiI*<sup>#</sup> | SAUSA300_1661 |  | tRNA | s<sup>4</sup>U8 |
| *dusB2* | SAUSA300_0089 |  | tRNA | D17, D20, D20a |
| *trmK* | SAUSA300_1520 |  | tRNA | m<sup>1</sup>A22 |
| *yjbO* | SAUSA300_0909 | NE822 | tRNA | ψ32 |
| *tadA* | SAUSA300_0543 | NE802 | tRNA | I34 |
| *tilS*<sup>#</sup> | SAUSA300_0487 |  | tRNA | k<sup>2</sup>C34 |
| *trmL* | SAUSA300_1805 | NE136 | tRNA | Cm34, Um34 |
| *tmcAL* | SAUSA300_1025 |  | tRNA | ac<sup>4</sup>C34 |
| *trhP1* | SAUSA300_1569 |  | tRNA | ho<sup>5</sup>U34 pathway |
| *thrP2*<sup>#</sup> | SAUSA300_1570 |  | tRNA | ho<sup>5</sup>U pathway |
| *trhO* | SAUSA300_2622 |  | tRNA | ho<sup>5</sup>U pathway |
| *trmR* | SAUSA300_1571 |  | tRNA | mo<sup>5</sup>U34 |
| *mnmA* | SAUSA300_1578 |  | tRNA | (c)mnm<sup>5</sup>s<sup>2</sup>U34 pathway |
| *yrvO*<sup>#</sup> | SAUSA300_1579 |  | tRNA | (c)mnm<sup>5</sup>s<sup>2</sup>U34 pathway |
| *mnmE* | SAUSA300_2646 |  | tRNA | (c)mnm<sup>5</sup>(s<sup>2</sup>)U34 pathway |
| *mnmG* | SAUSA300_2645 |  | tRNA | (c)mnm<sup>5</sup>(s<sup>2</sup>)U34 pathway |
| *mnmL*<sup>#</sup> | SAUSA300_1706 |  | tRNA | (c)mnm<sup>5</sup>(s<sup>2</sup>)U34 pathway |
| *mnmM* | SAUSA300_1707 | NE51 | tRNA | (c)mnm<sup>5</sup>(s<sup>2</sup>)U34 pathway |
| *folE2* | SAUSA300_0551 |  | tRNA | Q34 pathway |
| *queD* | SAUSA300_0696 |  | tRNA | Q34 pathway |
| *queE* | SAUSA300_0695 |  | tRNA | Q34 pathway |
| *queC* | SAUSA300_0697 |  | tRNA | Q34 pathway |
| *queF* | SAUSA300_0713 | NE1383 | tRNA | Q34 pathway |
| *tgt* | SAUSA300_1595 | NE1885 | tRNA | Q34 pathway |
| *queA* | SAUSA300_1596 |  | tRNA | Q34 pathway |
| *queG* | SAUSA300_1806 |  | tRNA | Q34 |
| *queH*<sup>#</sup> | SAUSA300_2528 |  | tRNA | Q34 |
| *trmD* | SAUSA300_1133 |  | tRNA | m<sup>1</sup>G37 |
| *trmM*<sup>#</sup> | SAUSA300_0464 |  | tRNA | m<sup>6</sup>A37 |
| *tsaC* | SAUSA300_2070 |  | tRNA | t<sup>6</sup>A37 |
| *tsaD* | SAUSA300_2002 |  | tRNA | t<sup>6</sup>A37 |
| *tsaB* | SAUSA300_2004 |  | tRNA | t<sup>6</sup>A37 |
| *tsaE* | SAUSA300_2005 |  | tRNA | t<sup>6</sup>A37 |
| *tcdA* | SAUSA300_1585 |  | tRNA | ct<sup>6</sup>A37 |
| *mtaB* | SAUSA300_1536 | NE1145 | tRNA | ms<sup>2</sup>t<sup>6</sup>A37 |
| *miaA* | SAUSA300_1195 | NE897 | tRNA | i<sup>6</sup>A37 |
| *miaB* | SAUSA300_1185 | NE1662 | tRNA | ms<sup>2</sup>i<sup>6</sup>A37 |
| *truA* | SAUSA300_2173 | NE223 | tRNA | ψ39 |
| *trmB* | SAUSA300_1694 | NE1672 | tRNA | m<sup>7</sup>G46 |
| *trmFO* | SAUSA300_1144 | NE1677 | tRNA | m<sup>5</sup>U54 |
| *truB* | SAUSA300_1164 | NE740 | tRNA | ψ55 |
| rRNA 16S |  |  |  |  |
| *rsuA* | SAUSA300_1699 | NE729 | rRNA 16S | ψ516 |
| *rsmG* | SAUSA300_2644 |  | rRNA 16S | m<sup>7</sup>G527 |
| *rsmD* | SAUSA300_1023 | NE1337 | rRNA 16S | m<sup>2</sup>G966 |
| *rsmB* | SAUSA300_1110 | NE924 | rRNA 16S | m<sup>5</sup>C967 |
| *rsmCII* | SAUSA300_0526 |  | rRNA 16S | m<sup>2</sup>G1050 |
| *rsmI* | SAUSA300_0466 | NE196 | rRNA 16S | m<sup>4</sup>Cm1402 |
| *rsmH*<sup>#</sup> | SAUSA300_1073 |  | rRNA 16S | m<sup>4</sup>Cm1402 |
| *rsmE* | SAUSA300_1537 | NE534 | rRNA 16S | m<sup>3</sup>U1498 |
| *rsmA* | SAUSA300_0470 |  | rRNA 16S | m<sub>2</sub><sup>6</sup>A1518, m<sub>2</sub><sup>6</sup>A1519 |
| rRNA 23S |  |  |  |  |
| *rlmCD* | SAUSA300_1878 |  | rRNA 23S | m<sup>5</sup>U747, m<sup>5</sup>U1939 |
| *rluD* | SAUSA300_1090 |  | rRNA 23S | Ψ1911, Ψ1915, Ψ1917 |
| *rlmH* | SAUSA300_0026 |  | rRNA 23S | m<sup>3</sup>Ψ1915 |
| *rlmB* | SAUSA300_0517 | NE936 | rRNA 23S | Gm2251 |
| *rlmL* | SAUSA300_1336 | NE1691 | rRNA 23S | m<sup>2</sup>G2445 |
| unknown |  |  | rRNA 23S | D2449 |
| *rluG* | SAUSA300_1800 |  | rRNA 23S | ψ2493 |
| *rlmN* | SAUSA300_1111 | NE1638 | rRNA 23S | m<sup>2</sup>A2503 |
| *rlmP* | SAUSA300_1036 |  | rRNA 23S | Gm2553 |
| *rlmQ* | SAUSA300_0981 | NE1546 | rRNA 23S | m<sup>7</sup>G2574 |
| *rluB* | SAUSA300_1443 | NE1244 | rRNA 23S | ψ2605 |

We examined the presence and sequence conservation of the 52 initial selected loci in 301 clinical *S. aureus* isolates from well-characterized cohorts of patients with pneumonia, bacteremia, endocarditis or cutaneous infections (28–30). The collection encompassed 30 clonal complexes and therefore represented substantial *S. aureus* genomic diversity. Each locus in the USA300_FPR3757 reference strain was compared with the corresponding sequences in the 301 genomes using different thresholds for sequence identity and alignment coverage (**Fig. 1**). At a minimum sequence-identity threshold of 95% (in green), most loci were detected in at least 298 of the 301 isolates. The principal exceptions were *rsmA* (261 strains) and *rluB* (276 strains) and the accessory antibiotic-resistance determinant *ermC* (15 strains) genes. Nevertheless, when the sequence-identity threshold was reduced to 90% (data not shown), *rsmA* and *rluB* were detected in all the 301 strains, indicating slightly higher sequence diversity at these loci rather than their absence from a subset of strains. By contrast, the acquired rRNA methylase gene *ermC* remained uncommon, being detected in no more than 16 isolates irrespective of the threshold applied. This limited distribution is consistent with the localization of *ermC* on mobile genetic elements associated with macrolide resistance (31, 32). Varying the minimum alignment-coverage threshold from 50% to 90% had little effect on the number of loci detected, indicating that most identified homologues covered at least 90% of the corresponding reference sequence. Thus, most loci in the original candidate set were broadly distributed and conserved across the clinical strain collection.

**Fig 1.**
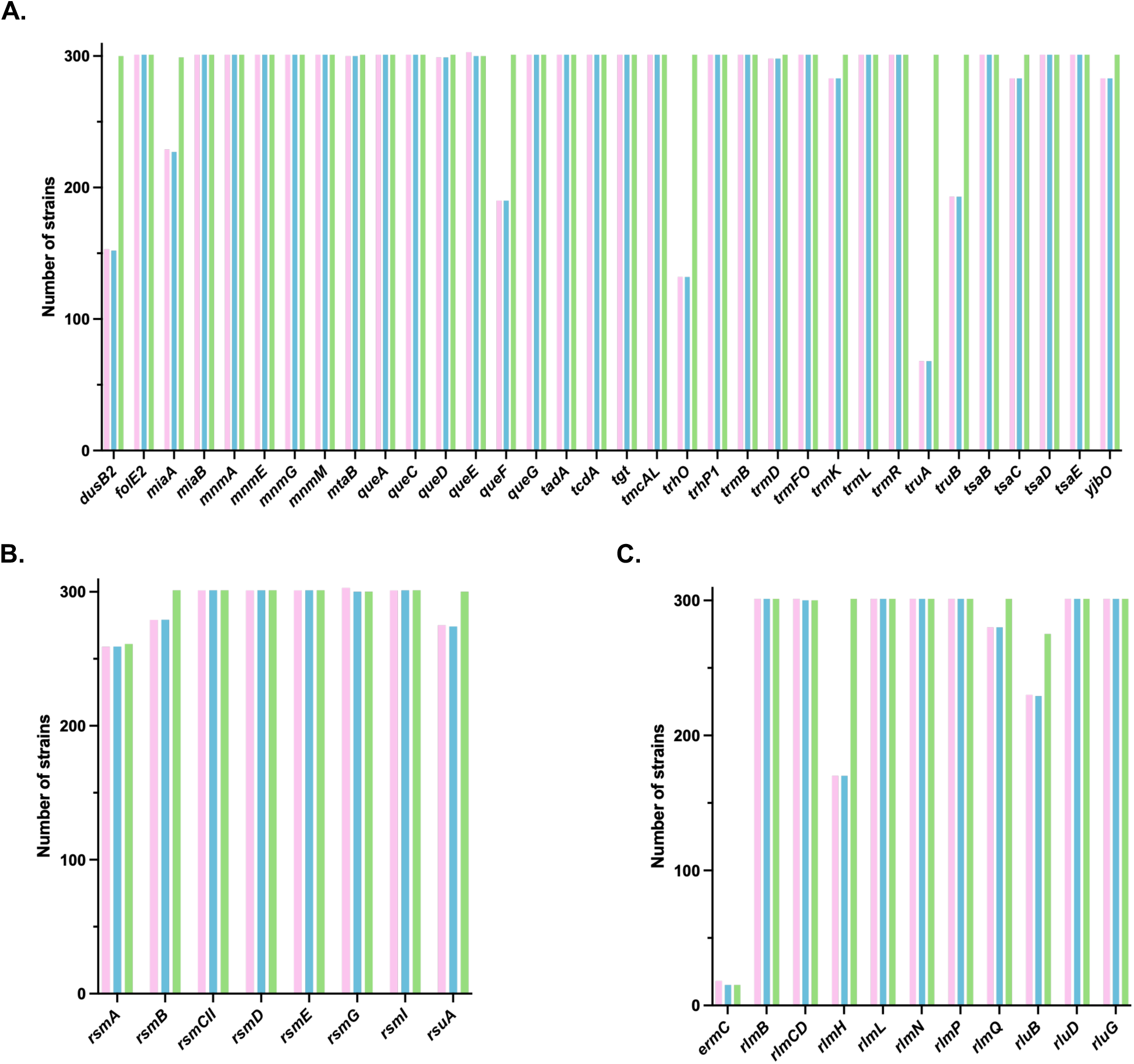
Presence of tRNA- and rRNA-modifying enzyme genes in *S. aureus* clinical strains. Genes coding for modifying enzymes targeting tRNA (A), 16S rRNA (B) and 23S rRNA (C) were searched in the genomes of 301 clinical strains by blasting the sequences from the reference strain USA300_FPR3757. Different filters were used to characterize the gene presence, using sequence homology and match length as parameters. The following thresholds were applied: in pink (match length ≥ 50% and homology ≥ 98%), in blue (match length ≥ 90% and homology ≥ 98%), and in green (match length ≥ 90% and homology ≥ 95%).

To assess conservation at protein-sequence level, we calculated a mutation rate for each candidate protein by dividing the number of observed amino-acid substitutions by the protein length (**Fig. 2**). For comparison, we analyzed three proteins expected to be highly conserved, GyrB, Hu and RplS, as well as the more variable coagulase protein CoA. The three conserved reference proteins had a mean mutation rate of 0.0008, compared with 0.1777 for CoA. Most proteins encoded by the candidate RNA-modification loci displayed similarly low values, ranging from 0.0004 to 0.0095. TrmK, RsuA and YjbO showed somewhat greater sequence variation, with values of 0.0152, 0.0138 and 0.0114, respectively, but remained substantially less variable than CoA.

**Fig 2.**
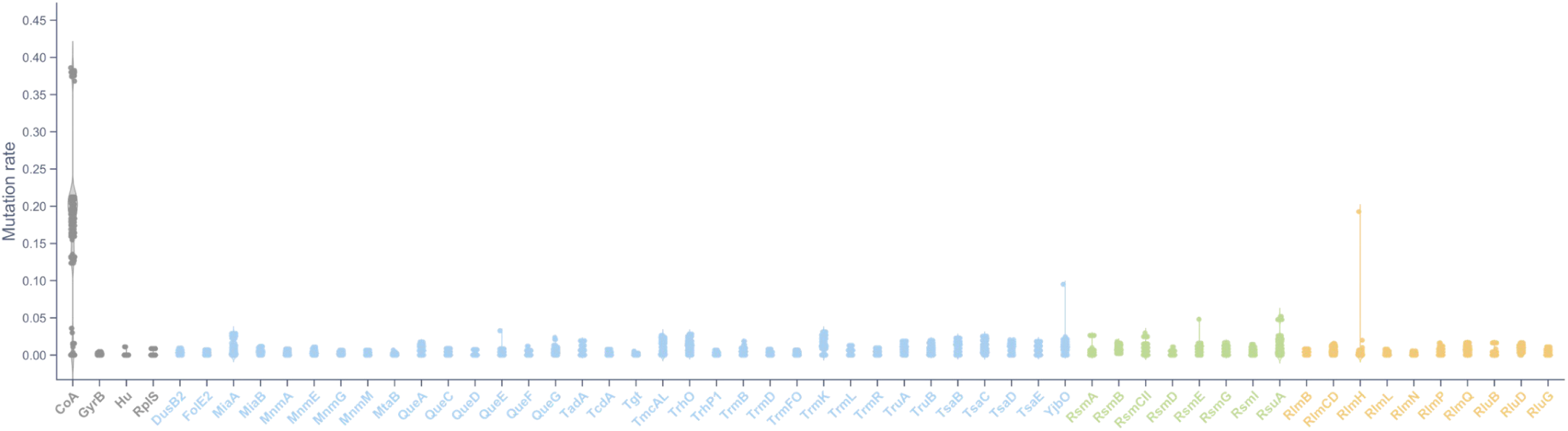
Protein sequences conservation of tRNA- and rRNA-modifying enzyme in *S. aureus* clinical strains. Mutation rates of protein sequences of tRNA (blue), 16S rRNA (green), and 23S rRNA (yellow) modifying enzymes, as well as control proteins (grey) from the 301 clinical strains were calculated by dividing the number of amino acid mutations by the sequence length.

Together, these analyses show that most loci are widely distributed and display limited sequence variation across genetically diverse clinical *S. aureus* isolates. They therefore define a broadly conserved repertoire of RNA-modification pathways for subsequent phenotypic analysis.

### Impact of RNA-modifying enzyme inactivation on bacterial growth

To investigate phenotypes associated with disruption of RNA-modifying enzyme genes in *S. aureus*, we selected strains from the Nebraska Transposon Mutant Library, which was generated in the USA300 JE2 background (33). Among the loci included in the genomic screen, 26 were represented by available transposon mutants and were selected for phenotypic analysis (**Table 1**). The position of each transposon insertion within the corresponding coding sequence was confirmed by sequencing. The growth, stress-associated phenotypes and virulence factors production of these mutants were compared with those of the parental J2 strain (WT).

We first examined growth under four culture conditions: in a nutrient-rich medium (BHI) at 30°C, 37°C or 42°C, and in RPMI cell culture medium at 37°C (**Fig. 3**). For each growth curve, we determined the doubling time during exponential growth and the final optical density at 600 nm (OD₆₀₀). None of the tested mutants showed a significant difference from the WT strain in either parameter under the BHI conditions examined (**Fig. 3A-F**). In RPMI at 37°C, no significant differences in exponential-phase doubling time were detected (**Fig. 3G**), but several mutants reached a significantly lower final OD_600_ than the WT strain (**Fig. 3H**). The mean final OD_600_ values of the *queF*^-^, *miaB*^-^, *trmFO*^-^, *trmB*^-^ and *rlmQ*^-^ transposon mutants were 0.32, 0.26, 0.16, 0.21 and 0.22, respectively, compared with 0.40 for the WT strain. Thus, these mutants displayed reductions in final OD₆₀₀ ranging from approximately 20% to 60%, without detectable changes in exponential growth rate. These results identify medium-dependent phenotypes associated with inactivation of several candidate RNA-modifying enzyme genes and suggest that these mutants have reduced culture yield or altered stationary-phase behavior in RPMI.

**Fig 3.**
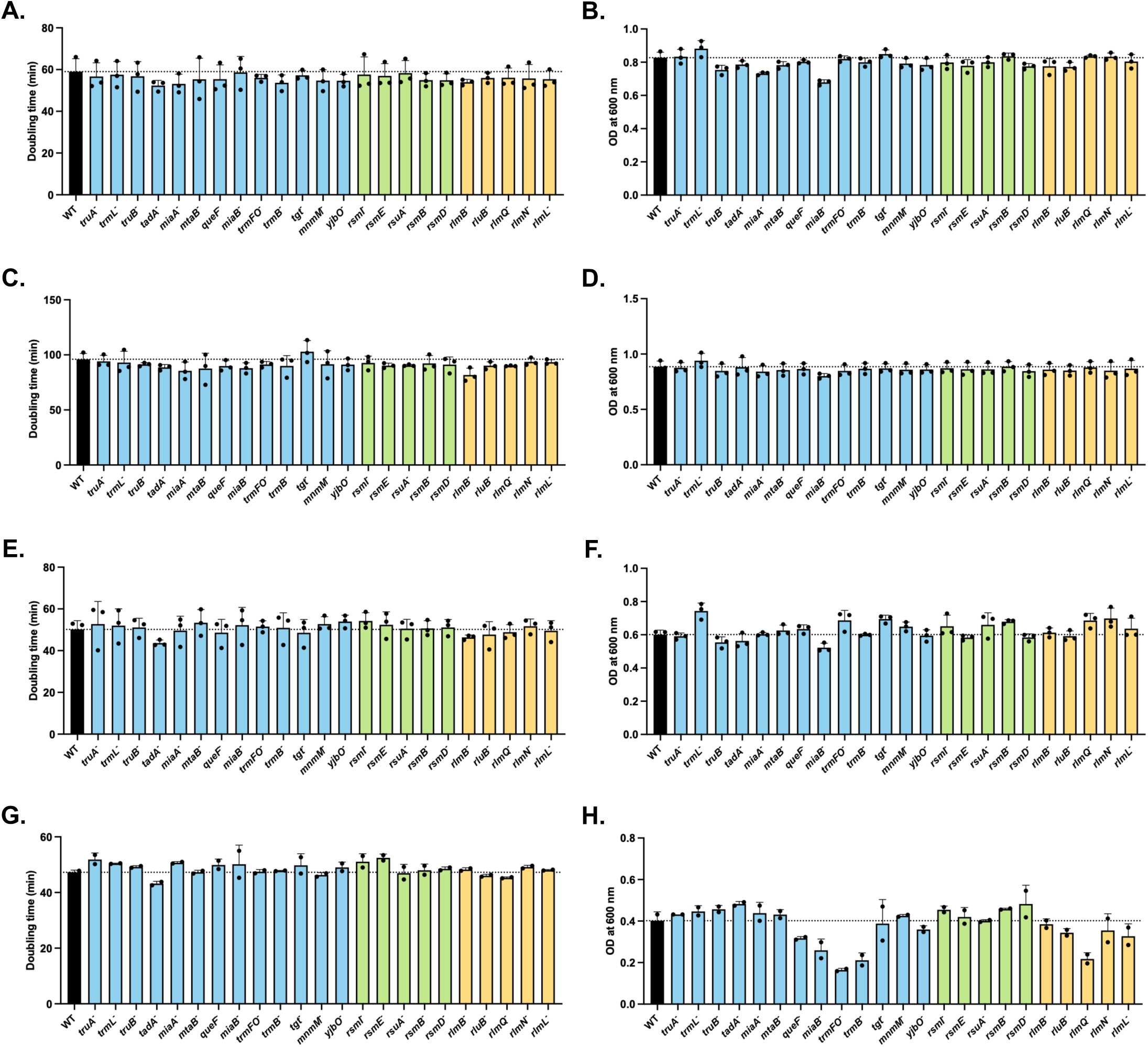
Bacterial growth of RNA-modifying enzyme transposon mutants. Bacterial growth was followed using a TECAN plate reader for 23 hours : in BHI medium at 37°C (A-B), 30°C (C-D) or 42°C (E-F), and in RPMI medium at 37°C (G-H), n=2, 3. Doubling time during the exponential growth phase (A, C, E et G) and final OD_600nm_ (B, D, F H) were analyzed. Mutant strains are colored according to the RNA type targeted by the inactivated enzyme: tRNA (blue), 16S rRNA (green) and 23S rRNA (yellow).

### Impact of RNA-modifying enzyme inactivation on antibiotic stress response

As antibiotic resistance contributes to *S. aureus* pathogenicity, we first determined the resistance profile using antibiotic susceptibility testing (AST) with Sensititre plates (**Fig. 4**). Overall, inactivation of RNA-modifying enzymes had no impact on antibiotic resistance as the transposon mutants displayed profiles similar to that of the WT strain. The only exception was the *rlmQ*^-^ mutant, which had an increased susceptibility to B-lactams (oxacillin and cefoxitin). It exhibited a cefoxitin minimal inhibitory concentration (MIC) of 4 µg/mL compared with 16 µg/mL for the WT strain, and an oxacillin MIC below 0.25 µg/mL compared with greater than 8 µg/mL for the WT strain.

**Figure 4.**
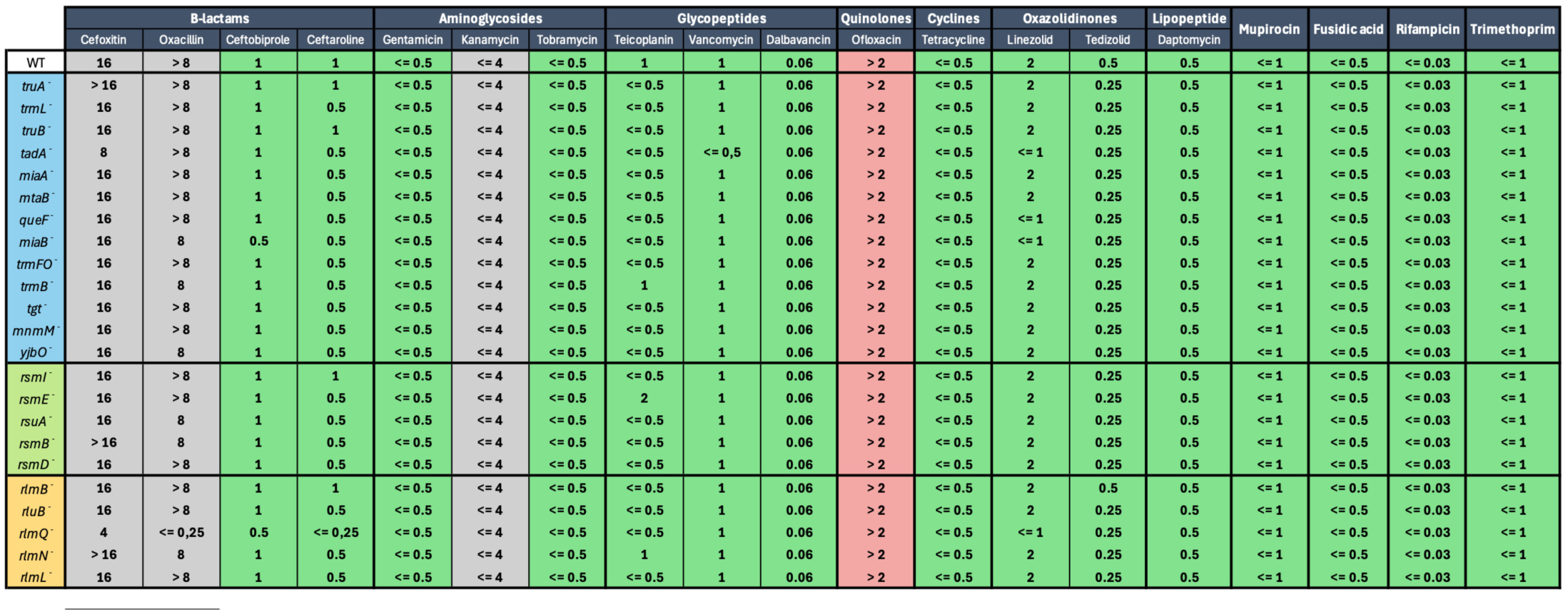
Resistance profiles of RNA-modifying enzyme transposon mutants. MIC has been determined using Sensititre plate after 24 hours of incubation in MH medium at 37°C, n=3. Cells background are colored according to resistance profile (sensitive in green, resistant in red and non-determined in grey).

Despite the limited changes in resistance profiles, we further sought to investigate bacterial growth and survival under antibiotic stress. Given the role of RNA modifications on translation, two bacteriostatic antibiotics targeting protein synthesis were selected: linezolid which inhibits the formation of the initiation complex by binding to the 50S subunit of the ribosome, and tetracycline which prevents tRNA binding by binding to the 30S subunit of the ribosome. Because the following experiments were conducted in BHI medium (rather than in MH medium as for Sensititre plates), MIC values were first determined in these conditions. No significant differences in MIC values were observed between the WT and mutant strains (**Table S1** - MIC was 1 or 2 µg/mL for linezolid and 0.25 µg/mL for tetracycline).

To evaluate the response to antibiotic stress of the different strains, cultures were grown in presence of sub-inhibitory antibiotic concentrations, defined as 0.25 times the MIC (**Fig. 5**). In the presence of linezolid, transposon mutants inactivated for tRNA-modifying enzymes (in blue) appeared to growth less efficiently than the WT strain, especially *truA*^-^ and *truB*^-^ mutants after 4 hours of culture (respectively 4.4 x 10^7^ and 6.7 x 10^7^ CFU/mL vs 1.5 x 10^8^ CFU/mL for WT) (**Fig. 5A**), as well as *truB*^-^ and *miaB*^-^ mutants after 8 hours of culture with respectively 6.8 x 10^8^ and 6.9 x 10^8^ CFU/mL compared with 1.1 x 10^9^ CFU/mL for the WT strain (**Fig. 5C**). A similar trend was observed in presence of tetracycline at 8 hours of culture, although these differences were not statistically significant (**Fig. 5D**).

**Fig 5.**
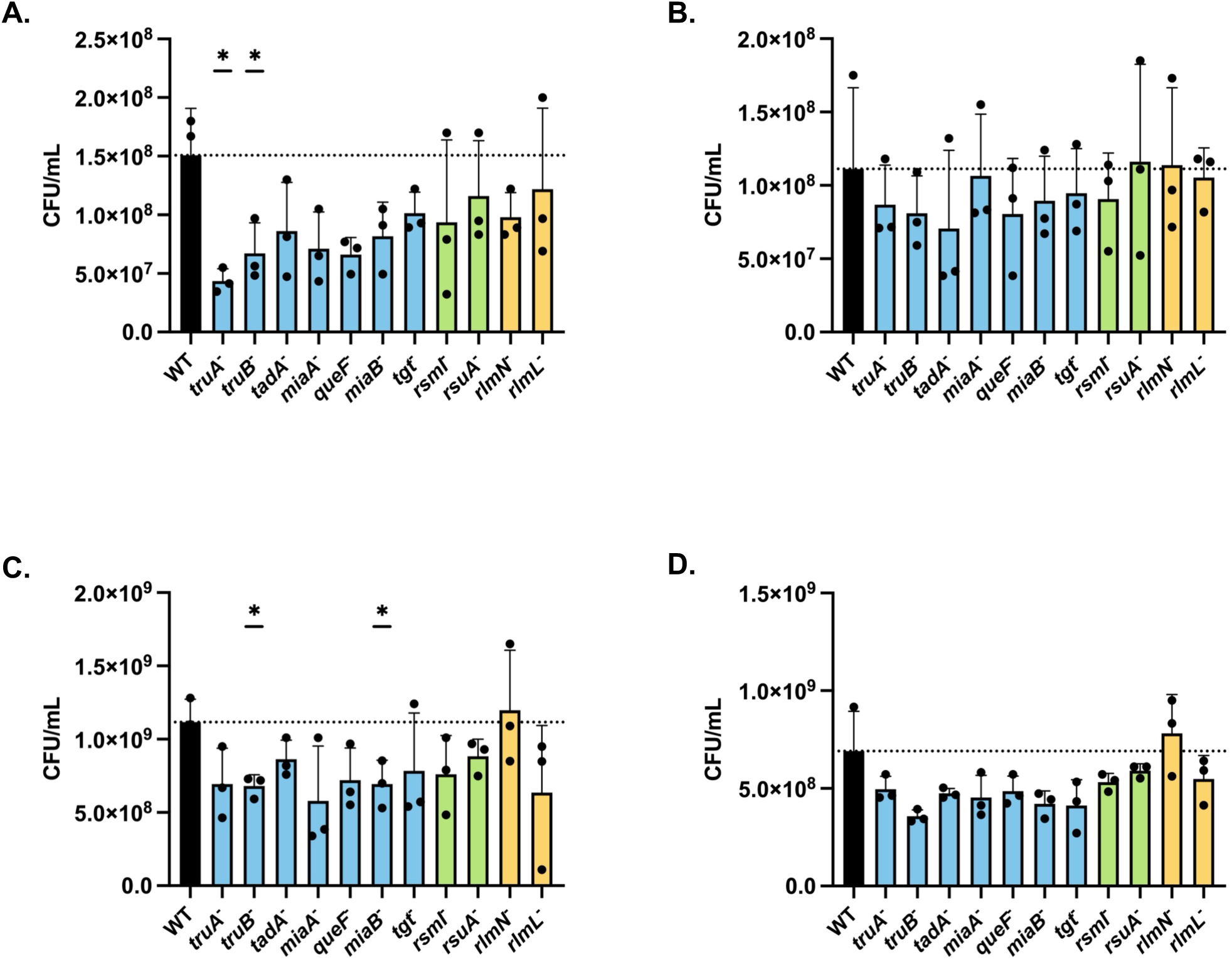
Growth of RNA-modifying enzyme transposon mutants in presence of antibiotic at sub-inhibitory concentrations. Bacterial growth in the presence of antibiotic at 0.25X MIC was monitored by counting CFU on TSA plates after 4 hours (A-B) and 8 hours (C-D) of culture, in presence of linezolid (A, C) or tetracycline (B, D), n=3. Statistical analysis were performed with Welch’s t test (A, C) or Mann-Whitney tests (B,D): (∗) P < 0.05, non-significative data are not shown.

Finally, we investigated bacterial survival during prolonged exposure of linezolid or tetracycline at 75 times the MIC (**Fig. 6**). For both antibiotics, the *truA*^-^, *miaA*^-^, and *miaB*^-^ mutants showed reduced survival compared with the WT strain at multiple time points. After 24 hours of exposure to linezolid (**Fig. 6C**), the survival rate was 1.7% for the WT strain, compared with 0.24% for *truA*, 0.26% for *miaA*^-^, and 0.09% for *miaB*^-^. Similarly, after 24 hours exposure to tetracycline (**Fig. 6F**), the survival rate was 0.87% for the WT strain compared with 0.44% for *truA*^-^, 0.39% for *miaA*^-^, and 0.41% for *miaB*^-^. Distinct antibiotic-specific phenotypes were also observed as the survival rate of certain transposon mutants appeared to be specifically affected by only one antibiotic. For example, the *truB*^-^ mutant showed reduced survival in the presence of linezolid with an average 2.1-fold less survival than the WT (**Fig. 6A–C**), whereas its survival rate remained similar to that of the WT strain in the presence of tetracycline (**Fig. 6D–F**). Conversely, the *tadA*^-^ mutant displayed no difference in survival when exposed to linezolid but showed approximately 2.8-fold lower survival than the WT strain during tetracycline exposure. These results indicate that inactivation of some RNA-modifying enzyme genes alters the response of *S. aureus* to ribosome-targeting antibiotics.

**Fig 6.**
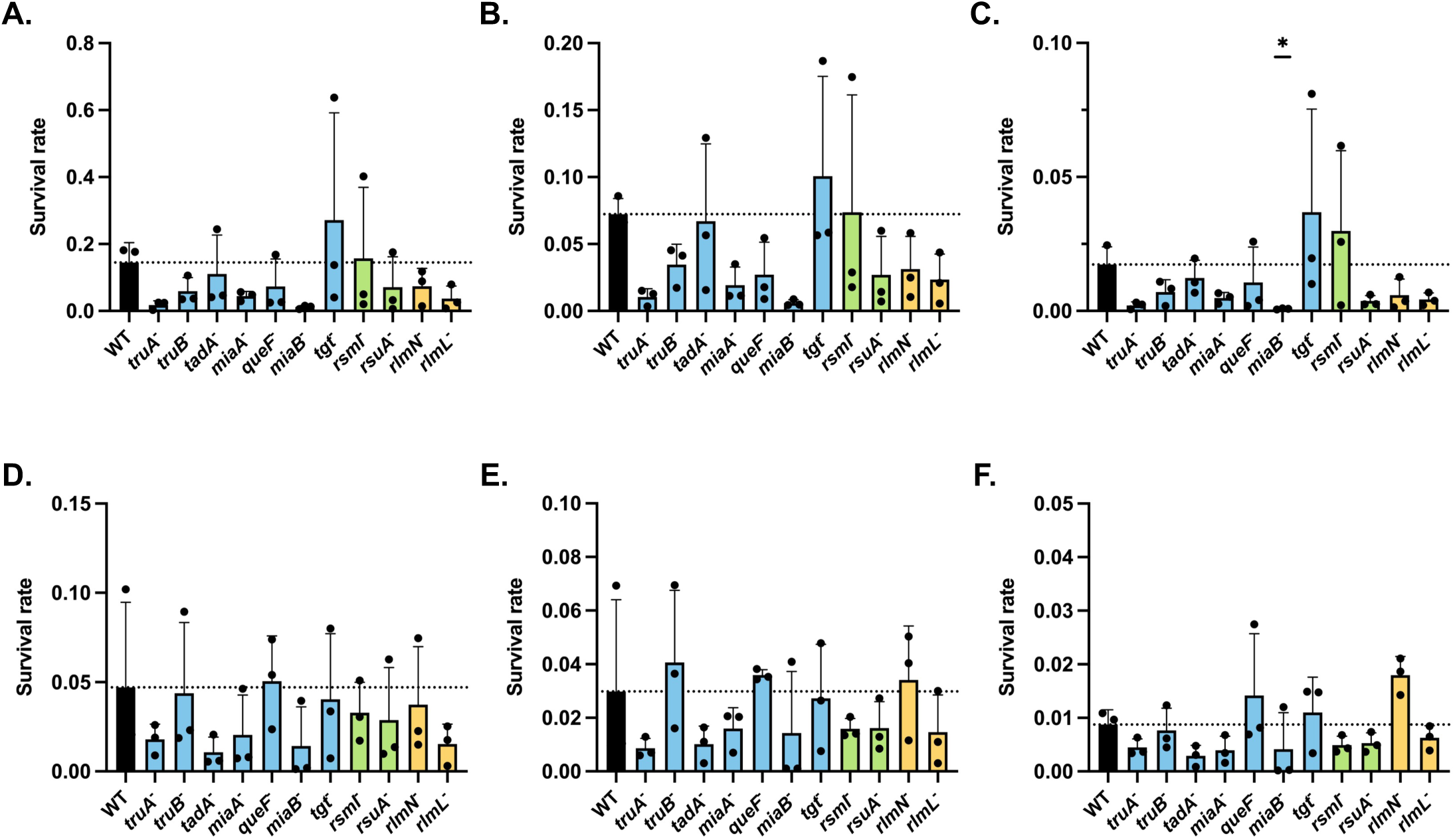
Survival of RNA-modifying enzyme transposon mutants during prolonged exposure to high antibiotic concentrations. Survival of bacterial strains exposed to antibiotic at 75X MIC was quantified by counting CFU on TSA plates after 4 hours (A, D), 8 hours (B, E) and 24 hours (C, F) of culture in presence of linezolid (A-C) or tetracycline (D-F). Survival rate was calculated relative to the initial inoculum set to 1, n=3. Statistical analysis were performed with Mann-Whitney tests (A, B, D-F) or Welch’s t test (C): (∗) P < 0.05, non-significative data are not shown.

### Impact of RNA-modifying enzyme inactivation on oxidative stress response

During infection, *S. aureus* is exposed to oxidative stress, primarily generated by the host phagocytes through the production of reactive oxygen species (ROS). To test the contribution of RNA-modifying enzyme to oxidative stress response, bacterial cultures were challenged with methyl-viologen after 2 hours of culture, and bacterial survival was then determined by CFU counting after 22 hours of culture. Among the transposon mutants tested, only the *miaB*^-^ mutant showed a clear and significant decrease in its survival rate, with 30% survival after 24 hours of culture compared with 450% for the WT strain (**Fig. 7A**).

**Fig 7.**
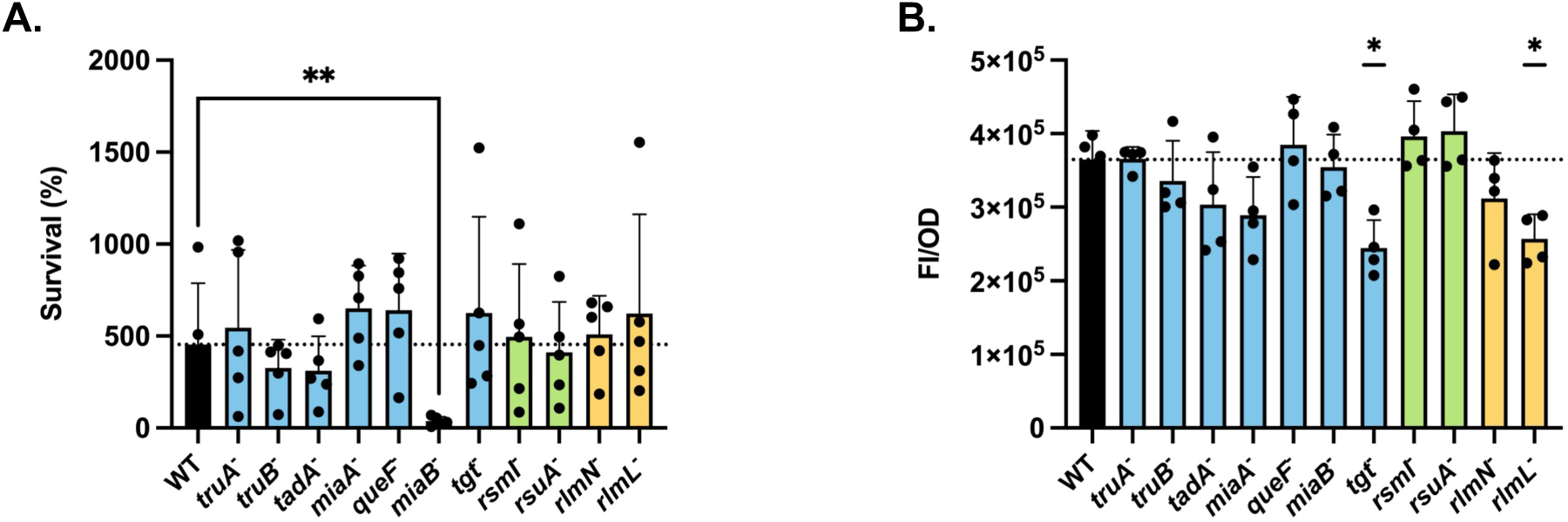
Tolerance of RNA-modifying enzyme transposon mutants to oxidative stress. (A) Bacterial viability after exposure to methyl viologen (10mM) was quantified by counting CFU on TSA plates after 22 hours of culture, n=5. (B) Endogenous ROS levels (Reactive Oxygen Species) after 6 hours of culture in BHI were measured by fluorescence intensity of the 2′,7′-dichlorofluorescein diacetate probe and normalized to OD_600nm_, n=4. Statistical analysis were performed with Mann-Whitney tests: (∗) P < 0.05, (∗∗) P < 0.01, non-significative data are not shown.

To investigate whether this phenotype was associated with altered ROS elimination, ROS endogenous levels were quantified. No difference was observed between the *miaB*^-^ mutant and the WT strain after 6 hours of culture, with ROS levels of 3.64 x 10⁵ IF/DO and 3.54 x 10⁵ IF/DO, respectively (**Fig. 7B**). Thus, these results indicate that the reduced survival of the *miaB*^-^ mutant under oxidative stress is not associated with an impaired ability to eliminate ROS in these conditions. In contrast, the *tgt*^-^ and *rlmL*^-^ mutants displayed lower endogenous ROS levels than the WT strain, reaching of 2.45 x 10⁵ IF/DO and 2.57 x 10⁵ IF/DO, respectively, after 6 hours of culture. However, this difference did not result in increased resistance to oxidative stress, as both mutants showed survival rates comparable to that of the WT strain following methyl-viologen exposure (**Fig. 7A**).

### Impact of RNA-modifying enzyme inactivation on cellular infection

The pathogenicity of *S. aureus* depends in part on its ability to adhere, invade, and survive within host cells. These key steps were examined upon infection of murine macrophages (RAW) by the different transposon mutants and the WT strain. Several mutants displayed altered interactions with macrophages (**Fig. 8**). The *rlmL* mutant exhibited significantly reduced adhesion (32%) compared to the WT strain (80%) (**Fig. 8A**). Similar reductions were observed for the *miaB*^-^ and *tgt*^-^ mutants with 40% and 36% of adhesion, respectively. The *miaB*^-^, *tadA*^-^ and *rlmN*^-^ mutants also showed reduced internalization (40%, 35%, and 37%, respectively) compared with 65% for the WT strain (**Fig. 8B**). In contrast, no significant differences were detected for intracellular survival among the strains tested (**Fig. 8C**). These results indicate that inactivation of RNA-modifying enzyme genes can affect early stages of macrophage interactions, including adhesion and internalization, without measurably altering intracellular survival under the conditions tested.

**Fig 8.**
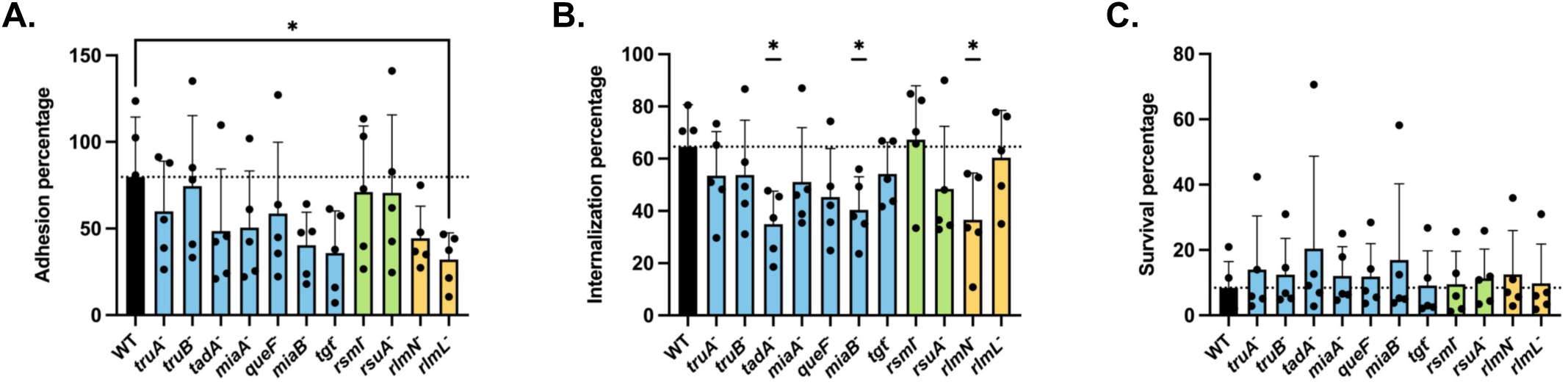
Adhesion, internalization and intracellular survival of RNA-modifying enzyme transposon mutants *in vitro*. Infection of RAW murine macrophages with bacterial strains at MOI 10, n=5. (A) Adhesion percentage corresponds to the proportion of adhered bacteria 2 hours post-infection, relative to the inoculum. (B) Internalization percentage corresponds to the proportion of internalized bacteria 3 hours post-infection, relative to adhered bacteria. (C) Survival percentage corresponds to the proportion of survival bacteria 24 hours post-infection, relative to the internalized bacteria. Statistical analysis were performed with Mann-Whitney tests: (∗) P < 0.05, non-significative data are not shown.

### Impact of RNA-modifying enzyme inactivation on toxin production

The production of major pore-forming toxins HlgC (**Fig. 9A**), PVL (**Fig. 9B**), and Hla (**Fig. 9C**), which contribute to *S. aureus* virulence, was quantified by ELISA after 18 hours of culture in CCY medium. In addition, Hla activity was functionally assessed by measuring hemolysis of rabbit red blood cells (**Fig. 9D**). The results suggest that RNA-modifying enzymes influence toxin production as several transposon mutants displayed different production profiles compared to the WT strain. The *trmL*^-^ mutant, for instance, produced significantly lower levels of both HlgC (55 ng/mL) and Hla (9.4 µg/mL) than the WT strain (126 ng/mL of HlgC and 13.9 µg/mL of Hla). Conversely, transposon insertion in the *tadA* gene notably increased the production of PVL (248 µg/mL compared with 188 µg/mL for the WT) and Hla (25.4 µg/mL compared with 9 µg/mL for the WT). In addition, the *miaB*^-^ mutant displayed distinct effects depending on the toxin, with a reduced PVL production (92 µg/mL) and an increased Hla production (19.2 µg/mL). Additional toxin-specific phenotypes were also observed. This is the case for the RlmN enzyme, whose inactivation reduced HlgC production by a factor of two compared with the WT strain (66 ng/mL for the *rlmN*^-^ mutant), for MiaA whose inactivation reduced PVL production (129 µg/mL vs 188 µg/mL for the WT), and for YjbO whose inactivation decreased Hla production (9.8 µg/mL vs 13.9 µg/mL for the WT). Together, these results indicate that inactivation of specific RNA-modifying enzymes differentially affects production of major *S. aureus* toxins.

**Fig 9.**
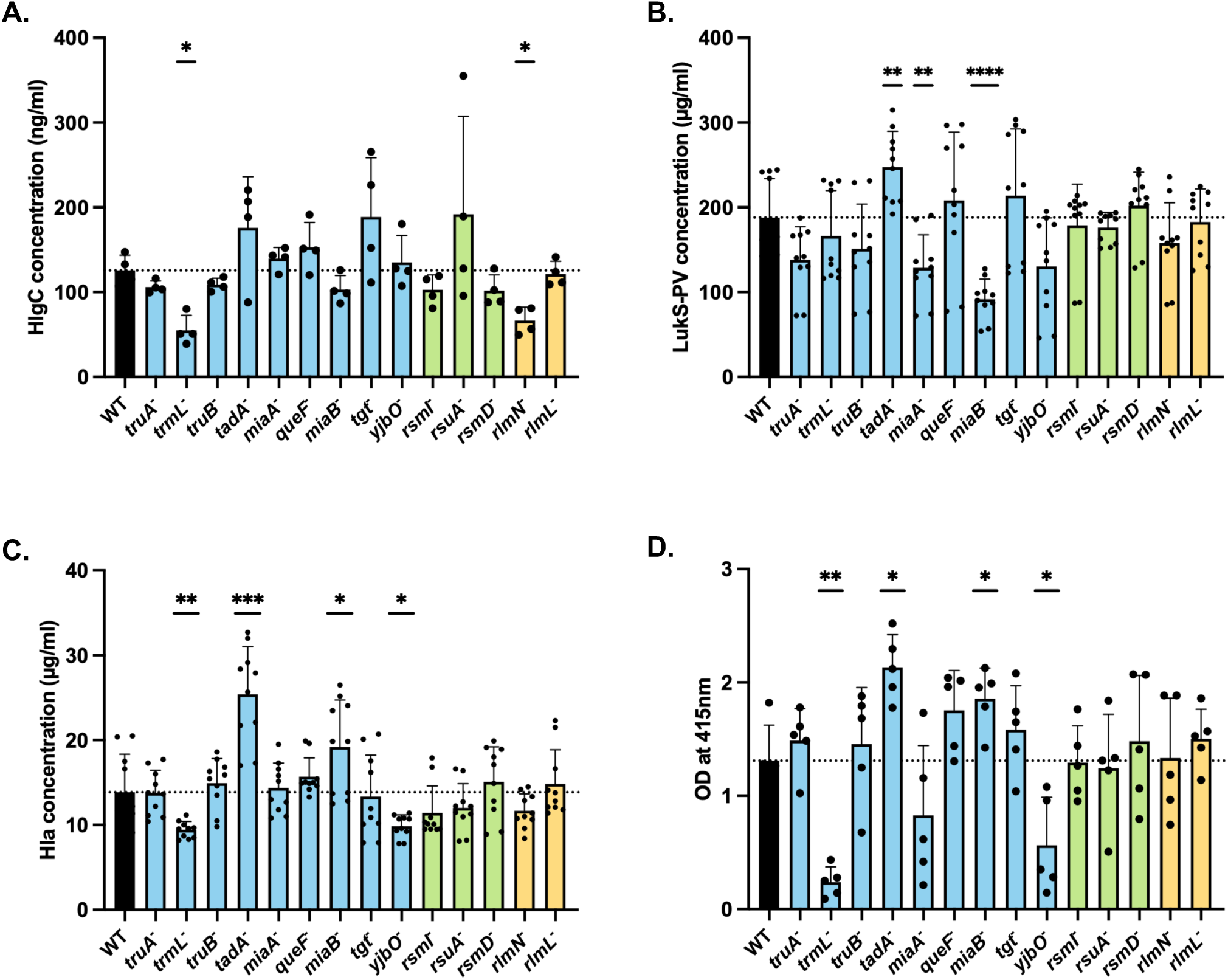
Toxins production and hemolytic activity of RNA-modifying enzyme transposon mutants. (A-C) Quantification of HlgC (A), PVL (B) and Hla (C) in bacterial supernatants from 18 hours of culture in CCY by ELISA assay, n= 4 à 10. (D) Evaluation of hemolytic activity of bacterial supernatants on rabbit red blood cells, measured by OD_415nm_, n=5. Statistical analysis were performed with Mann-Whitney tests: (∗) P < 0.05, (∗∗) P < 0.01, (∗∗∗) P < 0.001, (∗∗∗∗) P < 0.0001, non-significative data are not shown.

Finally, it is important to note that the variations in Hla production were consistent with the results obtained in the hemolysis test (**Fig. 9D**). The *tadA*^-^ and *miaB*^-^ mutants, which produced higher levels of Hla than the WT, also exhibited increased hemolysis activity, with an OD_415nm_ of 2.13 and 1.86, respectively, compared with 1.31 for the WT strain. Conversely, the *trmL*^-^ and *yjbO*^-^ mutants, whose Hla production were lower than that of the WT, showed significantly reduced hemolysis activity (0.14 and 0.28, respectively).

### Impact of RNA-modifying enzyme inactivation on the production of virulence factors

Because several RNA-modifying enzyme transposon mutants displayed altered toxin production, we expanded our analysis to include a broader range of virulence-associated proteins using a semi-quantitative targeted mass spectrometry method previously described (**Fig. 10**) (34). Several transposon insertions in RNA-modifying enzyme genes had opposite effects on different virulence factors: the *tadA*^-^ mutant produced half the amount of the SspP protease compared with the WT strain (ratio of 0.6), while producing higher levels of toxins (Hla = 1.7, Hld = 1.5, PSM-α = 1.6, LukSF = 1.5), as well as the immune evasion protein Chp (ratio of 2.3). Similarly, the *miaB*^-^ mutant produced half the amount of the SspA protease and the Nuc nuclease (ratios of 0.6) compared with the WT strain, while producing two-fold more the SplB protease, PSM-α and Hla toxins, as well as the Chp factor (ratios of 1.9).

**Fig 10.**
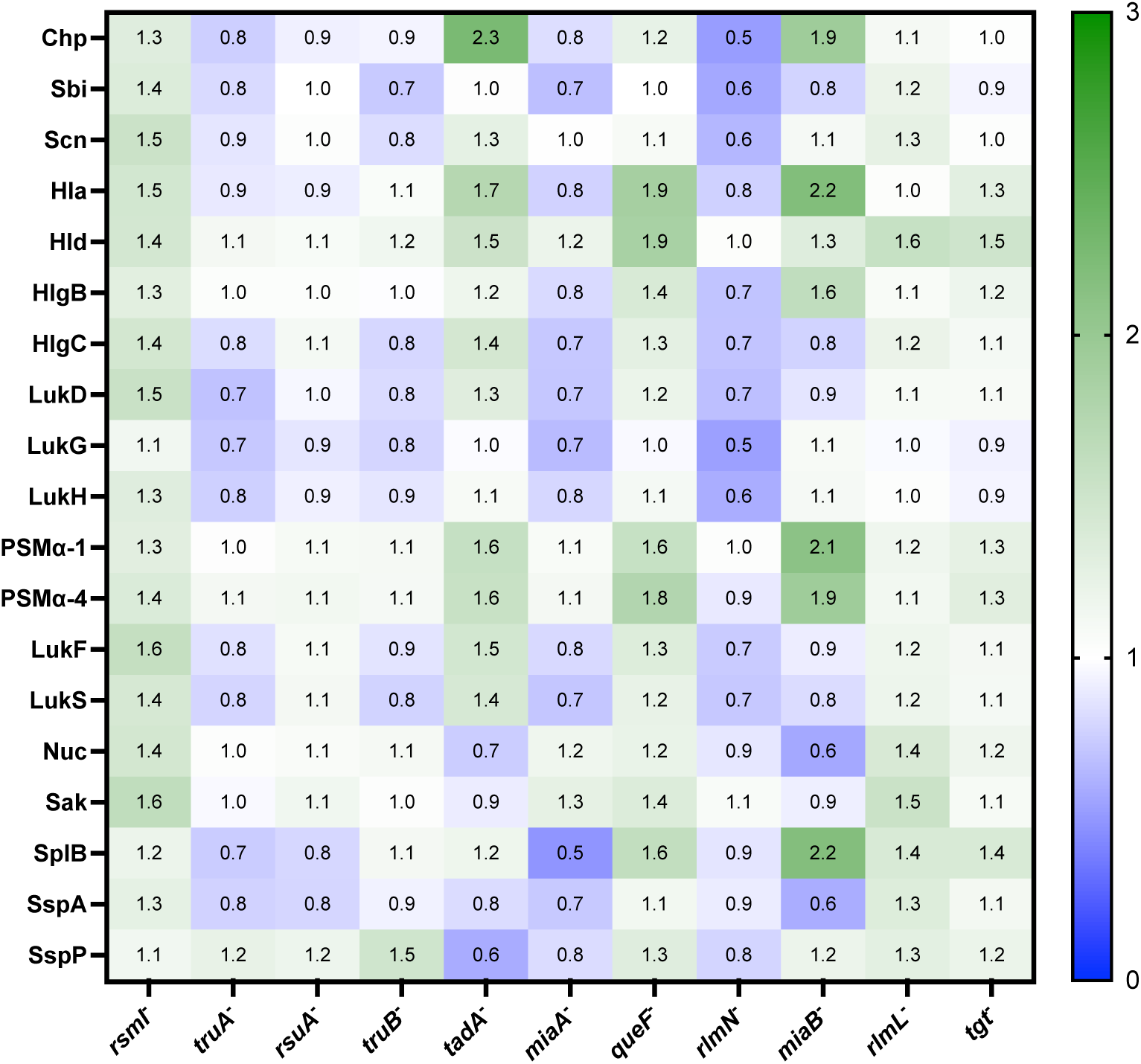
Production of virulence factors by RNA-modifying enzyme transposon mutants. Relative amount of 19 virulence factors were quantified by targeted mass spectrometry and normalized to ribosomal proteins. Relative amounts measured in mutant strains were compared to the WT strain using a ration. A value of 1 indicates no variation in the relative production between mutant and WT strains, n=3.

Other transposon mutants exhibited a protein expression profile that was uniformly different from that of the WT: for instance, *rsmI*^-^ and *queF*^-^ mutants produced slightly more of the different factors compared with the WT strain. Conversely*, rlmN*^-^ and *miaA*^-^ mutants showed a general decrease in the amount of virulence factors. In contrast, the other mutants did not show any significant change in virulence factor production, as *truA*^-^, *rsuA*^-^, *truB*^-^, *rlmL*^-^, and *tgt* mutants exhibited ratios close to 1 for almost all the virulence factors tested, indicating a production profile similar to that of the WT strain.

## DISCUSSION

In this study, we investigated the conservation and functional contribution of predicted tRNA- and rRNA-modifying enzymes in *S. aureus*. Genomic analysis demonstrated that most candidate loci belong to the core genome and are highly conserved among a diverse collection of clinical strains, defining a broadly conserved repertoire of putative RNA-modification pathways in *S. aureus*. As these predicted modifying enzymes primarily target tRNA and rRNA, two central molecules of the translation process, it may explain their high degree of conservation. However, it is important to note that the transposon mutant strains used are fully viable suggesting that these enzymes are not essential. Given the high number of modifications present on tRNAs and rRNAs, it is likely that these modifications fulfill complementary or even redundant functions, allowing some to compensate for the absence of others. In this context, it might be interesting to study mutants inactivated for multiple enzymes.

Importantly, phenotypic characterization of selected RNA-modifying enzymes mutants revealed that the transposon insertions did not affect growth under optimal laboratory conditions. Most mutants grew similarly to the WT strain in a rich medium, suggesting that these RNA modifications are not essential for growth in favorable environments. In contrast, several mutants displayed reduced final growth in RPMI medium, which mimics more nutrient limited conditions. This observation suggests that RNA-modifying enzymes may become important under conditions of environmental or host-associated stresses, potentially by optimizing protein production and metabolic adaption.

To support this hypothesis, our results highlight a potential role for RNA modifications in antibiotic stress response. Although most mutants showed no major changes in minimal inhibitory concentrations, several displayed altered growth or survival during exposure to sub-inhibitory or high concentrations of antibiotics targeting translation. These results suggest that RNA modifications do not directly determine antibiotic resistance levels but may influence bacterial fitness during antibiotic stress and, in some cases, survival during prolonged antibiotic exposure. In particular, the *truA*^-^, *truB*^-^ *miaA*^-^ and *miaB*^-^ mutants showed a reduced growth under antibiotic stress. Consistent with these findings, inactivation of *truA*, *truB* and *miaB* was previously reported to reduce *Vibrio cholerae* fitness during exposure to sub-inhibitory concentrations of tobramycin, gentamycin, as well as ciprofloxacin for the *truA* and *miaB* mutants, and chloramphenicol for the *truA* mutant (20).

Similar to antibiotic stress, the response to oxidative stress further supports a condition specific contribution of RNA-modifying enzymes to bacterial adaptation. The *miaB*^-^ transposon mutant displayed a strong reduction in survival following oxidative stress, although its endogenous ROS levels were not increased relative to the WT strain. Thus, the reduced survival of the mutant does not appear to result from increased basal ROS accumulation but may instead reflect defects in stress-responsive translation, protein homeostasis or metabolic adaptation. MiaB is a radical-SAM methylthiotransferase containing two Fe–S clusters, and its activity may therefore be sensitive to cellular redox conditions, as has been described for other enzymes (35, 36). However, whether MiaB itself functions as a redox sensor in *S. aureus* remains to be established. Interestingly, we recently detected ms²A37 in *S. aureus* tRNA^Leu^(UAA), suggesting that MiaB can introduce the methylthio group directly onto unmodified A37, independently of prior MiaA-mediated isopentenylation (14). Loss of this unusual MiaB-dependent modification could affect the translation of proteins required for oxidative-stress adaptation. The phenotype observed here contrast with those reported in an extraintestinal pathogenic *E. coli* strain, in which deletion of *miaA*, but not that of *miaB*, impaired growth in the presence of the superoxide-generating agent methyl viologen (37), suggesting that the functions of these enzymes in oxidative-stress adaptation vary between bacterial species. Finally, some mutants displayed reduced intracellular endogenous ROS levels without improved survival after oxidative stress exposure, indicating that ROS accumulation alone does not fully explain the phenotypes observed in presence of oxidative stress.

Host cell infection assays revealed that several transposon mutants were affected in adhesion or internalization into macrophages, while intracellular survival remained mostly unaffected. These results suggest that RNA modifications may influence early stages of host-cell interactions rather than to bacterial intracellular persistence. This could be linked to altered expression of surface proteins, adhesins or regulators involved in host-cell interactions. Notably, the reduced adhesion observed for *rlmL*^-^ mutant and the reduced internalization for *miaB*^-^, *rlmN*^-^ and *tadA*^-^ mutants indicates that distinct RNA modification pathways may differentially affect specific steps of cellular infection.

As observed for cell infection, the impact of RNA modification pathways on toxin and virulence factors production was highly dependent on the enzyme inactivated. Some transposon insertions decreased the production of specific toxins, whereas others increased toxin levels or differentially alter other virulence factors production, highlighting the diversity of phenotypic outcomes associated with individual RNA modifications. For example, *tadA* inactivation was associated with increased production of several toxins, whereas *rlmN^-^*and *miaA*^-^ mutants showed a more general decrease in virulence factor production. In addition, the *miaB*^-^ mutant displayed opposite effects depending on the virulence factor studied. These results illustrate that RNA-modifying enzymes may not only enhance or repress virulence in a global way but rather reshape the balance of virulence factors production.

The phenotypes identified in this study may be explained, at least in part, by the established roles of tRNA and rRNA modifications in modulating translation efficiency by influencing RNA stability, decoding accuracy, codon usage bias and ribosome activity (7, 15). In this context, the expression of specific virulence-associated genes may be particularly sensitive to the loss of specific RNA modifications, especially if their translation depends on particular codon usage or requires optimal ribosomal activity. Therefore, the loss of specific tRNA or rRNA modifications could selectively alter the production of certain proteins without affecting overall protein production and bacterial growth. Therefore, future experiments combining ribosome profiling, proteomics and codon usage analysis would help determine whether the observed phenotypes result from specific effects on translation. Such post-transcriptional regulation may allow *S. aureus* to adjust protein synthesis quickly and specifically in response to changing conditions and RNA modifications thus appear to be a subtle way of driving bacterial virulence program and stress responses.

A major finding of this study is that tRNA- and rRNA-modifying enzymes do not appear to act as global regulators of *S. aureus* virulence, unlike major regulatory systems such as Agr or SaeRS which coordinate broad transcriptional responses (38, 39). The impacts observed here were highly specific to the enzymes inactivated, the phenotypes tested and the conditions. Rather than supporting a single global regulatory role of tRNA and rRNA modifications, this highlights a set of candidate pathways contributing to the fine-tuning of bacterial adaptation rather than functioning as master regulators. Interestingly, it is already known that under non optimal conditions, many modifying enzymes and RNA modifications are important for bacterial physiology, particularly in the presence of environmental or host-associated stresses (40–43).

Among the RNA-modifying enzymes tested, MiaB inactivation was associated with the largest number of phenotypic alterations., including antibiotic and oxidative stress responses, host-cell interactions and toxin production. This suggests that the MiaB-dependent tRNA methylthiolation, primarily associated with ms²i⁶A37 formation, may have a broader role in staphylococcal physiology than some other tRNA- or rRNA-modification pathways studied here (44). However, the contribution of MiaB to bacterial adaptation and virulence remains poorly characterized, as only a limited number of studies exist. One described MiaB as positive regulator of the T3SS in *P. aeruginosa*, therefore promoting cytotoxicity in human lung epithelial cells (44) and another showed that MiaB is involved in metabolism and morphogenesis in *Streptomyces ghanaensis* as its inactivation reduces moenomycin production and alters cellular morphology (45). Therefore, further investigations on MiaB-dependent tRNA methylthiolation and its downstream consequences on gene expression will be interesting to understand the molecular basis of the multiple phenotypes observed in *S. aureus* in this work.

This study has several limitations. First, the phenotypic screening was performed using transposon mutants, and genetic complementation will be required to confirm that the observed phenotypes are directly attributable to the inactivation of the targeted RNA-modifying enzymes. Second, the corresponding RNA modifications were not directly quantified, and future studies should use dedicated approaches such as LC-MS/MS or RNA sequencing to confirm the loss of specific modifications. Third, most experiments were conducted in a single genetic background, whereas *S. aureus* clinical strains are highly diverse. Validation in additional genetic backgrounds and in relevant *in vivo* infection models will therefore be important to determine the broader significance of these findings.

Nevertheless, our results demonstrate that RNA-modification pathways contribute to the adaptation and virulence of *S. aureus* in a multifactorial aspect and condition-specific manner. It appears that RNA-modifying enzymes contribute to a context-dependent regulation of gene expression, enabling *S. aureus* to modulate virulence factors expression. tRNA and rRNA modifications thus constitute a post-transcriptional regulatory system, like other mechanisms such as regulatory RNAs, riboswitches, or two-component systems, that allow bacteria to respond to metabolic and environmental stimuli. This regulatory flexibility likely supports the temporal and spatial control of virulence factor expression during infection, thereby promoting bacterial adaptation and pathogenicity. These findings expand our understanding of post-transcriptional regulation in *S. aureus* and provide a basis for future studies establishing the mechanisms underlying the contribution of individual RNA-modifying enzymes and RNA modifications as modulators of bacterial pathogenicity.

## MATERIAL AND METHODS

### Bacterial strains

All transposon mutant strains used in the phenotypic screening are indicated in **Table 1** and were obtained from the Nebraska Transposon Mutant Library constructed in the USA300_JE2 background (33). Transposon insertion in the targeted gene has been verified by sequencing.

### Genomic analysis

Bioinformatics analysis was performed on genomes of 301 *S. aureus* clinical strains previously sequenced (28–30). First, the 52 candidate loci included in the original genomic screen were searched by BLAST in the 301 genomes using the USA300_FPR3757 sequences as references. Then, protein sequences of the RNA-modifying enzymes, as well as sequences of CoA (SAUSA300_0224), GyrB (SAUSA300_0005), Hu (SAUSA300_1362) and RplS (SAUSA300_1134) proteins from these 301 clinical strains, were aligned and compared to the protein sequences from USA300_FPR3757 strain using Clustal Omega version 1.2.4 (46). A mutation rate was calculated by dividing the number of mutated amino acids by the sequence length. Newly curated candidates added to **Table 1** after this analysis were not included in these comparative genomic analyses.

### Growth curve

Strains were cultured in BHI broth at 37°C overnight with shaking (200 rpm) and diluted at OD_600_ =0.03 in fresh medium. 96-well plate was seeded with 200 µL and incubated at 30°C, 37°C or 42°C in a Tecan plate reader for 24 hours. OD_600_ was measured every 15 min after agitation and the doubling time was calculated during the exponential growth phase.

### MIC determination

#### Sensititre plate

Strains were streaked on Tryptic Soy Agar (TSA) plates and incubated for 24 hours at 37°C. Bacterial suspensions were subsequently prepared at 0.5 McF in MH medium, and 50 µL were inoculated into Sensititre plates (Thermo Scientific). The plates were incubated during 24 hours at 37°C and the MICs were read using a Vizion plate reader (Thermo Scientific).

#### Microdilution

Strains were streaked on TSA agar plate and incubated for 24 hours at 37°C. Bacterial suspensions adjusted to 0.5 McF were prepared in BHI broth and diluted at 1:50. A U-bottom 96-well plate was seeded with 50 µL of BHI, 50 µL of diluted inoculum and 50 µL of antibiotic solution. A two-fold serial dilution of the antibiotic was realized across the columns. The plate was then incubated for 24 hours at 37°C, and the MIC was identified as the lowest concentration of antibiotic presenting no growth.

### Growth with antibiotic stress

Strains were cultured in BHI broth at 37°C overnight with shaking (200 rpm) and diluted at OD_600_ =0.05 in 20 mL of fresh medium with antibiotic. Bacterial suspensions were cultured during maximum 24 hours at 37°C with shaking (200 rpm). Samples were collected at different time points to measure OD_600_ and enumerate bacteria by colony counting on TSA plates.

### Survival under oxidative stress

Strains were cultured in BHI broth at 37°C overnight with shaking (200 rpm) and diluted at OD_600_ =0.1 in 20 mL of fresh medium and incubated for 2 hours at 37°C with shaking. Oxidative stress was then induced by adding methyl viologen at 10 mM. Samples were collected at different time points to measure OD_600_ and enumerate bacteria by colony counting on TSA plates.

### ROS quantification

Strains were cultured in BHI broth at 37°C overnight with shaking and diluted at OD_600_ =0.05 in 20 mL of fresh medium before incubation at 37°C with shaking. After 6 hours of culture, endogenous ROS production was quantified as previously described (47).

### *In vitro* infection

Murine macrophages (RAW) cell line was grown in DMEM GlutaMAX medium (Gibco) supplemented with 10% fetal bovine serum at 37°C and 5% CO_2_. 24-well tissue culture plates were seeded with 100 000 cells per well. After 24 hours of incubation, the cells were washed twice with 1 mL of Phosphate Buffered Saline (PBS, Gibco) and infected at a multiplicity of infection (MOI) of 10:1 with bacterial suspensions from 8-hour bacterial cultures and diluted in DMEM supplemented medium. After 2 hours at 37°C and 5% CO_2_, cells were washed with 1 mL of PBS and lysed by osmotic shock with 1 mL of ultra-pure water. The collected cell lysate was plated on TSA agar to determine the number of bacteria adhering to the cells. To determine the number of internalized bacteria, cells were infected for 2 hours, treated with 1 mL of lysostaphin at 10 µg/mL and gentamycin at 400 µg/mL (Sigma-Aldrich) for one hour before cell lysis to eliminate extracellular bacteria. The cell lysate was then plated on TSA agar. Intracellular survival was assessed at 24 hours post-infection, with antibiotic treatment (lysostaphin at 5 µg/mL and gentamycin at 200 µg/mL) maintained until the end of the experiment. The collected cell lysate after was plated on TSA agar to determine the number of surviving intracellular bacteria.

### Toxin quantifications by ELISA

Bacterial strains were grown overnight in 10 mL of CCY medium at 37°C with shaking (200 rpm) and then centrifuged during 10 minutes at 10 000 g to collect the supernatants. Quantification of HlgC, LukSF and Hla toxins were carried out by a sandwich ELISA assay using custom made antibody as previously described (48–50).

### Red blood cells lysis

Bacterial strains were cultured overnight in 10 mL of CCY medium at 37°C with shaking (200 rpm) and then centrifuged during 10 minutes at 10 000 g to collect the supernatants. Serial dilutions of the supernatants (from 1/2 to 1/2048) in PBS were prepared and 100 µL of each dilution were added to a U-bottom 96-well plate. Then, 50 µL of rabbit red blood cells (Atlantis) washed in PBS and resuspended at 10^8^ cells/mL were added. After 1 hour of incubation at 37°C, the plate was centrifuged and the optical density at 415 nm was measured to characterize the hemolytic activity of bacterial strains.

### Targeted proteomics

Strains were cultured in CCY broth at 37°C overnight with shaking (200 rpm) and diluted at OD_600_ =0.05 in 5 mL of fresh medium. After 8 hours of culture at 37°C with shaking, 1 mL of bacterial suspensions were collected. Sample preparations and proteomic analysis were realized as previously described (34).

## ACKNOWLEDGMENTS

This work was supported by the French National Research Agency ANR (SaRNAmod:ANR-21-CE12-0030-01 to FV; RHU IdBioriv: ANR-18-RHUS to FV), and by Fondation pour la Recherche Médicale FRM (FDT202404018387 to CT). The funders had no role in study design, data collection and analysis, decision to publish, or preparation of the manuscript.

## Figure legends

**Table S1.** MIC for linezolid and tetracycline in BHL. MIC has been determined by microdilution after 24 hours of incubations in BHI medium 37°C, n=4.

|  | Linezolid | Tetracycline |
|---|---|---|
| WT | 2 µg/mL | 0.25 µg/mL |
| *truA*<sup>-</sup> | 1 µg/mL | 0.25 µg/mL |
| *truB*<sup>-</sup> | 1 µg/mL | 0.25 µg/mL |
| *tadA*<sup>-</sup> | 1 µg/mL | 0.25 µg/mL |
| *miaA*<sup>-</sup> | 1 µg/mL | 0.25 µg/mL |
| *queF*<sup>-</sup> | 1 µg/mL | 0.25 µg/mL |
| *miaB*<sup>-</sup> | 2 µg/mL | 0.25 µg/mL |
| *tgt*<sup>-</sup> | 2 µg/mL | 0.25 µg/mL |
| *rsmI*<sup>-</sup> | 1 µg/mL | 0.25 µg/mL |
| *rsuA*<sup>-</sup> | 1 µg/mL | 0.25 µg/mL |
| *rlmN*<sup>-</sup> | 2 µg/mL | 0.25 µg/mL |
| *rlmL*<sup>-</sup> | 2 µg/mL | 0.25 µg/mL |

